# Aneuploidy and deregulated DNA damage response define haploinsufficiency in breast tissues of *BRCA2* mutation carriers

**DOI:** 10.1101/729301

**Authors:** Mihriban Karaayvaz, Rebecca E Silberman, Adam Langenbucher, Srinivas Vinod Saladi, Kenneth N Ross, Elena Zarcaro, Andrea Desmond, Murat Yildirim, Varunika Vivekanandan, Hiranmayi Ravichandran, Ravindra Mylavagnanam, Michelle C Specht, Sridhar Ramaswamy, Michael Lawrence, Angelika Amon, Leif W Ellisen

## Abstract

Women harboring heterozygous germline mutations of *BRCA2* have a 50-80% risk of developing breast cancer, yet the early pathogenesis of these cancers is poorly understood. We sought to reveal early steps in *BRCA2*-associated carcinogenesis through analysis of sorted cell populations from freshly-isolated, non-cancerous breast tissues among a cohort of *BRCA2* mutation carriers and matched controls. Single-cell whole-genome sequencing demonstrates that >25% of *BRCA2* carrier (*BRCA2*^*mut/+*^) luminal progenitor (LP) cells exhibit sub-chromosomal copy number variations (CNVs), which are rarely observed in non-carriers. Correspondingly, primary *BRCA2*^*mut/+*^ breast epithelia exhibit spontaneous and replication stress-induced DNA damage together with attenuated replication checkpoint and apoptotic responses, associated with an age-associated expansion of the LP compartment in human carrier tissues. These phenotypes are not associated with loss of wild-type *BRCA2*. Collectively, these findings provide evidence for *BRCA2* haploinsufficiency and associated DNA damage in vivo that precede histologic abnormalities. These results provide unanticipated opportunities for new cancer risk assessment and prevention strategies in high-risk patients.

## Introduction

Breast cancers arising in women who inherit heterozygous mutations in *BRCA2* are associated with a high prevalence of genomic alterations and aggressive clinical behavior (*1, 2*). Due to the high risk of these cancers in *BRCA2* mutation carriers, many such women elect to undergo bilateral mastectomy for breast cancer prevention. Yet despite the unmet need for more effective breast cancer prevention approaches in this setting, the stepwise evolution from an otherwise normal *BRCA2* heterozygous mutant (*BRCA2*^*mut/+*^) cell to an invasive malignancy has not been defined. Homozygous loss of *BRCA2* is embryonic lethal (*3-5*), and acute loss in cultured cells rapidly leads to DNA damage and growth arrest or cell death (*6-8*). These observations suggest a multi-step pathogenesis in which homozygous *BRCA2* loss is not the earliest genetic event, but rather that the wild-type *BRCA2* allele may remain intact as early genetic changes accumulate. Critically however, this scenario leaves unresolved the nature and enabling mechanism for early cancer evolution. Haploinsufficiency for *BRCA2* has been proposed as a possible driver of early pathogenesis, but direct evidence for such an effect in the normal human mammary gland is inconsistent. Furthermore, heterozygous genetically engineered mouse models (GEMMs) of *BRCA2* are not tumor prone and therefore represent a poor model of precancerous evolution in this setting (*3-5, 8, 9*). While the *BRCA1* tumor suppressor shares many of these features (*9, 10*), the pathogenesis of *BRCA1*-versus *BRCA2*-associated breast cancers may differ in important ways, as the former are primarily hormone receptor (HR) and HER2-negative tumors, while the latter are primarily HR-positive (*11*).

We sought to unveil the earliest steps in the pathogenesis of *BRCA2*-associated breast tumors through detailed analysis of histologically normal glands from women harboring germline deleterious mutations who elected to undergo bilateral prophylactic mastectomy. Genomic analysis of individual cells revealed frequent polyclonal chromosomal damage, which was most prevalent among the subset of epithelial cells that are the suspected cell-of-origin of these cancers. Corresponding defects in replication stress and DNA damage checkpoint responses in these same cells collectively define a previously unappreciated haploinsufficient phenotype for *BRCA2* in the human breast. The discovery of these precancerous hallmarks paves the way for improving clinical risk prediction and cancer prevention in this population.

## Results

### Single-cell whole genome analysis reveals sub-chromosomal aneuploidy in *BRCA2*^*mut/+*^ human primary breast epithelial cells

We carried out detailed analysis of histologically normal glands from *BRCA2* carriers who elected to undergo bilateral prophylactic mastectomy, using as controls tissues from age-matched women electing cosmetic breast surgery (Fig. 1A). We employed established markers to carry out flow cytometry-based isolation and sorting of the three major epithelial cell sub-populations: mature luminal (ML), luminal progenitor (LP) and basal epithelial cells (Fig. 1A). Notably, data from GEMMs and gene expression analyses of human tumors have suggested that the cell of origin of *BRCA1*-associated breast cancer is the LP cell (*12, 13*), while *BRCA2*-associated tumors may arise from an LP-related cell or a more mature luminal cell (*14*).

**Fig. 1.**
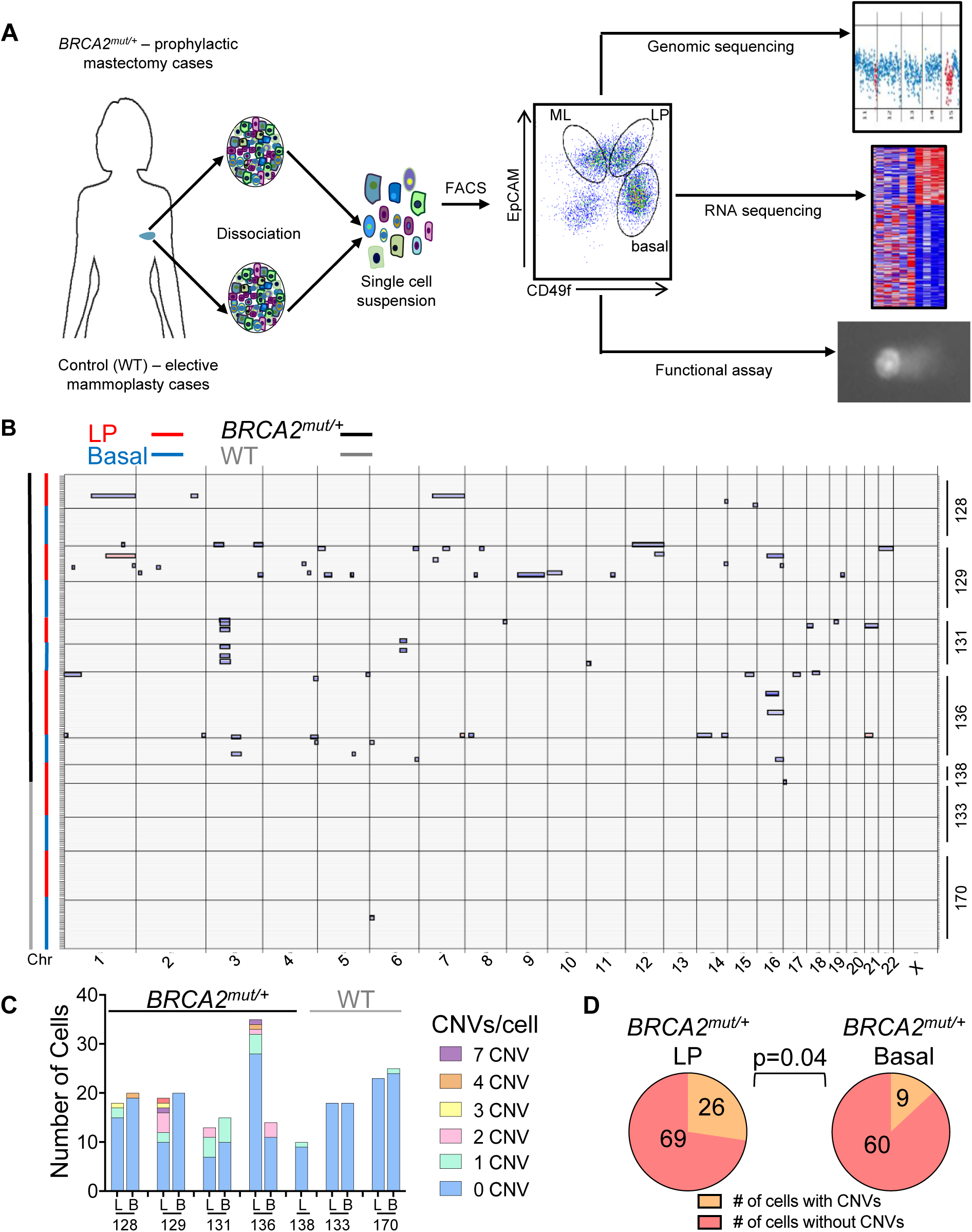
Single-cell whole genome analysis of *BRCA2*^*mut/+*^ human primary breast epithelial cells. **(A)** Workflow depicts dissociation and isolation of human breast epithelial cells from *BRCA2* carrier (*BRCA2*^*mut/+*^) prophylactic mastectomy and control (WT) elective mammoplasty cases for subsequent analyses as indicated. Dot plot at center shows representative flow cytometry sorting via CD49f and EpCAM of mature luminal (ML), luminal progenitor (LP), and basal epithelial cells. (A) Summary of single-cell whole-genome sequencing (WGS) analysis of flow-sorted, primary uncultured breast epithelial cells. Copy Number Variation (CNV) calls for individual cells (rows) across the genome (x-axis; Chr, chromosome) are shown, with gains and losses boxed. Cell types and genotypes are indicated at top left, and individual patient ID numbers are indicated at right. In total, 252 sequenced breast epithelial cells are depicted. **(C)** Bar chart depicting the prevalence of CNVs in LP (L) and basal (B) cells of *BRCA2* carrier and control (WT) patients. Color code depicts the number of CNVs identified per cell. **(D)** LP cells from *BRCA2* carriers are significantly more likely than basal cells to harbor CNVs. P-value by Chi-square test.

Among the earliest events in cancer evolution are thought to be polyclonal somatic genomic alterations. Accordingly, we looked for the presence of somatic CNVs at high-resolution through single-cell whole-genome sequencing (WGS) of uncultured, flow-sorted primary LP and basal epithelial cells from *BRCA2* carriers and controls. Low-coverage WGS provides sufficiently high resolution to identify sub-chromosomal CNVs as small as 10MB, and our methodology for single-cell whole-genome amplification and analysis has been previously validated (*15, 16*). We carried out WGS to an average depth between 0.1-0.05X, then used two independent algorithms (HMMcopy and DNAcopy) to assign and confirm copy number changes across the genome (*15, 16*). Previous studies employing this methodology have demonstrated that in unselected individuals the proportion of cells with any such CNVs is very low (<5% of cells) in normal epithelial and brain tissues (*16*). In contrast, among nearly 100 individual LP cells from a cohort of *BRCA2* carriers analyzed by WGS we observed that 27% demonstrated one or more CNVs of >10MB (Fig. 1, B to D). Applying this methodology to an equal number of basal breast epithelial cells from the same individuals also revealed a substantial excess of cells harboring CNVs (13%), although significantly less than the proportion of CNV-positive LP cells (p=0.04) (Fig. 1, B to D). By comparison, a parallel WGS analysis of sorted LP and basal cells from non-carriers revealed a single CNV in 90 cells (Fig. 1, B and C). As further validation of our sequencing and analysis pipelines we re-analyzed existing data from normal skin and brain cells sequenced on the same platform. The overall sequence quality was comparable between these cells and the breast epithelial cells, and we confirmed the low prevalence of CNV-positive cells in 142 skin and brain cells sequenced (fig. S1). Thus, breast epithelia and particularly LP cells from non-cancerous breast tissue of *BRCA2* carriers harbor frequent sub-chromosomal aneuploid events (Fig. 1D, fig. S2A).

One notable CNV we observed was duplication of the entire chromosome 1q arm, which is a common genomic abnormality in breast cancer (Fig. 2A) (*17*). The majority of the identified CNVs were sub-chromosomal haploid losses, consistent with the widespread pattern of losses observed in *BRCA2*-associated breast cancer (Fig. 2A and fig. S2B) (*1*). In some cases, identical losses were shared between multiple cells of the same patient, a finding which could conceivably correspond to early clonal evolution (Fig. 2B). Importantly, none of the losses in any cell involved the *BRCA2* locus on chromosome 13 (Fig. 1B and fig. S2, B to E). Prior analyses of germline *BRCA2*-associated breast cancers have demonstrated that most genetic loss-of-heterozygosity (LOH) events for *BRCA2* itself are >10MB and therefore would have been detected by our analysis (*18*). This observation suggests that the wild type *BRCA2* allele is intact in our cases, implying that accumulation of sub-chromosomal aneuploidy may be a haploinsufficient phenotype. To confirm the integrity of the wild-type allele we performed targeted PCR amplification of the locus surrounding the patient-specific *BRCA2* mutation from individual cells. Although efficiency for detection of either allele was relatively low, in no case did we identify a cell with a detectable mutant allele and no detectable wild-type allele (Fig. 2C). Taken together, these findings imply that sub-chromosomal aneuploidy is an early, haploinsufficient phenotype in *BRCA2*^*mut/+*^ breast epithelia.

**Fig. 2.**
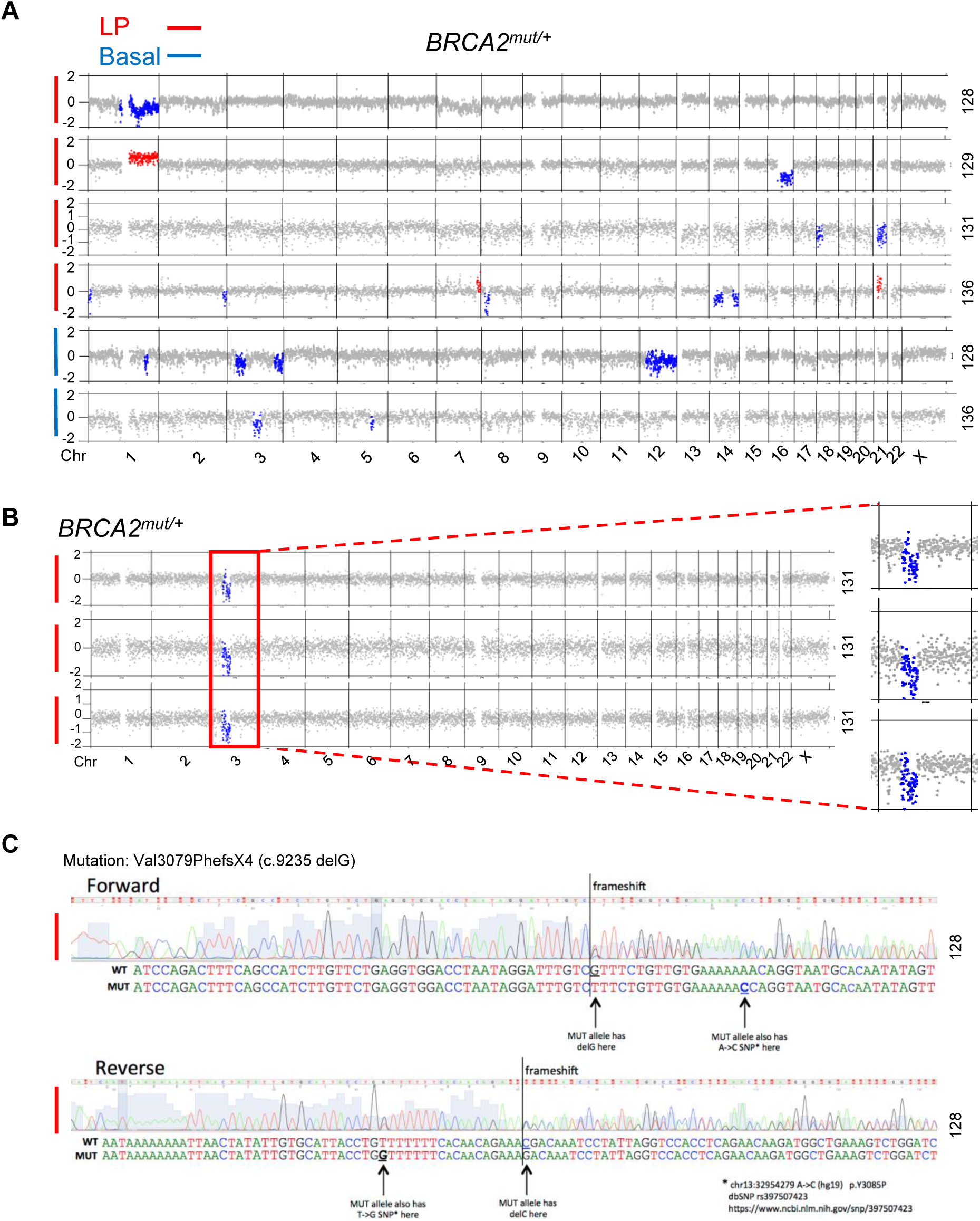
Polyclonal, sub-chromosomal aneuploidy is a hallmark of *BRCA2*^*mut/+*^ breast epithelial cells. **(A)** Representative segmentation plots of individual LP (n = 4) and basal (n = 2) cells harboring CNVs from four *BRCA2* mutation carriers. Y-axis depicts normalized WGS read counts across the genome (x-axis). Red dots indicate region of gain, blue dots indicate losses. Patient ID numbers are indicated at right. **(B)** Segmentation plots of 3 LP cells that share a clonal loss (red box) in a *BRCA2* carrier (131). Zoomed images of the clonal loss are shown at right. **(C)** Representative chromatograms from single-cell PCR-based Sanger sequencing of genomic DNA in a *BRCA2*^*mut/+*^ LP cell. The presence of a heterozygous SNP and the superimposition of sequences adjacent to the frameshift mutation suggest LOH has not occurred.

### *BRCA2*^*mut/+*^ primary cells exhibit DNA damage and a deregulated replication stress response

The presence of viable aneuploid cells in *BRCA2*^*mut/+*^tissues suggested ongoing DNA damage and a deregulated stress or damage response. Thus, we next used an independent method to directly assess DNA damage in single cells, the comet assay. This assay employs cells embedded in agarose that are lysed then subjected to electrophoresis, causing broken DNA structures to migrate toward the anode, thus forming a comet tail (*19*). We briefly cultured freshly-collected cells from *BRCA2* carriers or controls under ultra-low attachment conditions (48-72h) to select for epithelial progenitor cells prior to plating (*20*). Consistently, cells from *BRCA2* carriers demonstrated increased DNA breaks at baseline compared to controls (Fig. 3A). Additionally, inducing replication stress by treatment with hydroxyurea (HU) led to further increases in DNA damage in *BRCA2*^*mut/+*^ cells, potentially reflecting the established role of *BRCA2* in protection of stalled replication forks (Fig. 3A) (*21*).

**Fig. 3.**
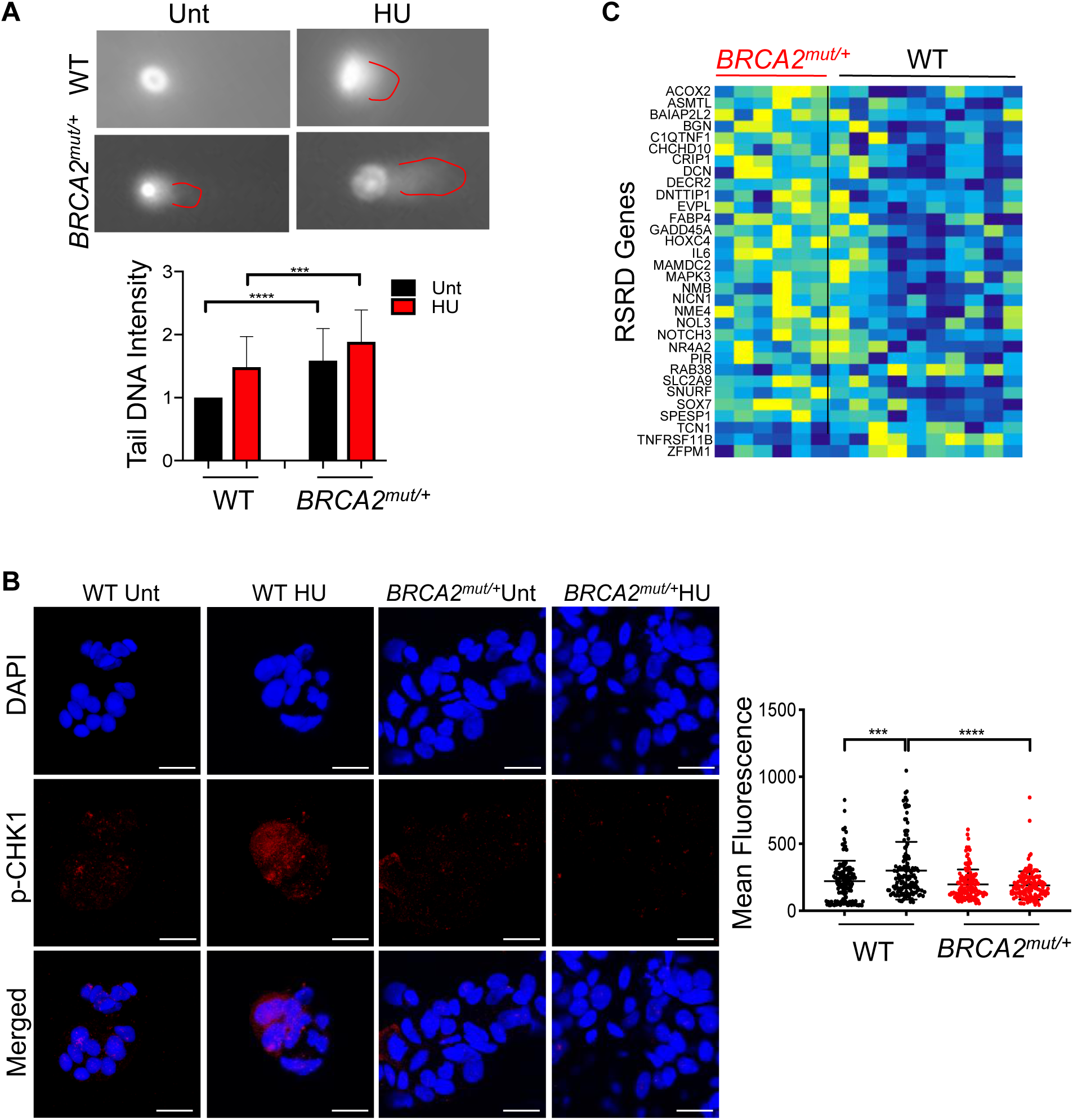
*BRCA2*^*mut/+*^ breast epithelial cells exhibit DNA damage and an impaired replication stress checkpoint response. **(A)** Representative images of comet assays performed on primary human breast epithelial cells isolated from control (WT) and *BRCA2*^*mut/+*^tissues. Red lines highlight “tail” of broken DNA. Graph below summarizes data from n = 3 patients per genotype, 50 cells per patient. Cells were either untreated (Unt) or treated with hydroxyurea (HU) for 4 h. Data are depicted as fold change in tail DNA intensity. p values by unpaired *t*-test. *** p < 0.001. Error bars indicate standard deviation. **(B)** Representative confocal immunofluorescence staining of primary breast epithelial cells for p-CHK1 (Ser317) shows increased nuclear staining following HU treatment only in control (WT) but not in *BRCA2*^*mut/+*^cells. Graph at right shows nuclear fluorescence of individual cells (dots) (n =4 patients for control, n = 3 for *BRCA2*^*mut/+*^; four fields counted per condition per patient). p values by unpaired *t*-test. *** p < 0.001, **** p < 0.0001. Horizontal lines indicate means and standard deviations. Scale bars represent 20 µm. **(C)** Heatmap of RNA-seq data from freshly-sorted cells shows differential expression of RSRD (replication stress response defect) genes (*24*) in *BRCA2*^*mut/+*^ LP cells (n = 7 patients) compared to control (WT) LP cells (n = 9 patients). Columns correspond to individual patients.

We then examined the response to this genomic stress by analyzing phosphorylation of CHK1, a central coordinator of the response to replication stress and DNA damage (*22, 23*). Cytospins of primary epithelial progenitor cultures prepared as above were stained for phosphorylated CHK1 at baseline or following 4 hours of exposure to HU. As anticipated, control primary epithelia exhibited increased CHK1 phosphorylation within 4 hours of HU treatment (Fig. 3B). In contrast, however, cells from *BRCA2* carriers exhibited a failure to active CHK1 in response to HU, despite normal levels of total CHK1 protein (Fig. 3B and fig. S3A). DNA sequencing of these cells revealed the presence of both wild-type and mutant *BRCA2* alleles (fig. S3B). These findings provide further support for a haploinsufficient phenotype of *BRCA2* in the response to genomic stress.

Because we observed a deregulated genomic stress response in vitro, we wanted to know whether this also occurs in vivo. Thus, we carried out RNA-seq analysis of freshly-sorted LP and basal epithelial cell populations from *BRCA2* carrier tissues or controls (Fig. 1A). Analysis of these data revealed enrichment in *BRCA2*^*mut/+*^ LP cells of an established signature reflecting a failure of the ATR/CHK1-mediated replication stress checkpoint in non-transformed mammary epithelial cells (Fig. 3C and fig. S3C) (*24*). This replication stress response deficiency (RSRD) signature is known to predict future cancer risk (*24*), and it contains some of the top most differentially expressed genes between *BRCA2*^*mut/+*^and control LP cells (Fig. 3C). Among these are genes of potential relevance to HR-positive breast cancer (which comprise 80% of *BRCA2*-associated breast cancers), including the estrogen receptor target gene *HOXC4* and the GATA transcription factor binding partner gene *ZFPM1* (Fig. 3C) (*25, 26*). Furthermore, evaluation of differentially expressed programs through Gene Set Enrichment Analysis revealed the highly significant deregulation of a radiation response signature in *BRCA2*^*mut/+*^ LP cells (fig. S3D) (*27*). Notably, the differential expression of this signature between *BRCA2*^*mut/+*^and control cells was far more significant within the LP compared to the basal population, in keeping with the more frequent occurrence of CNVs among LP cells (fig. S3D). Again consistent with haploinsufficiency for *BRCA2*, the RNA-seq data showed no evidence for exclusive expression of the mutant allele in *BRCA2*^*mut/+*^LP cells (fig. S3E). Thus, *BRCA2*^*mut/+*^ LP cells exhibit evidence of aberrant replication stress and DNA damage responses in vivo.

### *BRCA2*^*mut/+*^ LP cells show increased TP53 activity and decreased NF-kB/SASP pathway expression

We then turned to examine the downstream consequences of the DNA damage detected in LP cells of *BRCA2* carriers. A hallmark genetic event that cooperates with *BRCA2* deficiency in cancer pathogenesis is loss of *TP53*, suggesting that activation of TP53 may be an early barrier to malignant progression in this setting (*28*). We therefore hypothesized that the failed CHK1-dependent replication stress response we observed might ultimately lead to DNA double-strand breaks and thereby trigger TP53 activation through a CHK1-independent pathway (*29*). Indeed, recent studies suggest CHK1 is not required for TP53 activation in primary breast epithelial cells following DNA damage (*30*). RNA-seq analysis did suggest activation of TP53 in *BRCA2*^*mut/+*^ LP cells, evidenced by the increased expression of multiple direct TP53 target genes (Fig. 4A) (*31, 32*). This TP53 activation in vivo was associated with a strong transcriptional profile indicating suppression of NF-kB signaling, including numerous cytokine and inflammatory factors associated with the senescence-associated secretory phenotype (SASP) (Fig. 4, B to D). TP53 is known to suppress the NF-kB/SASP response (*33, 34*), and this effect is emerging as a relevant component of TP53-dependent tumor suppression given that accumulation of SASP-expressing cells is an established driver of tumorigenesis (*35*). We independently validated the corresponding alterations in NF-kB protein expression, demonstrating that the NFKB1 (p50) and NFKB2 (p52) subunits were expressed at lower levels in *BRCA2* carrier tissues compared to controls (Fig. 4C). Furthermore, knockdown of *BRCA2* in non-transformed mammary epithelial cells via lentiviral shRNA attenuated expression of the same cytokine and NF-kB targets genes that were downregulated in *BRCA2*^*mut/+*^progenitor cells in vivo (fig. S4A). Like the damage response signature (fig. S3D), deregulation of the SASP program was selective for LP cells in *BRCA2* carriers, as no significant suppression of SASP was observed in the corresponding basal epithelial cells of these same patients (fig. S4B). These results suggest that DNA damage and TP53 activation in *BRCA2*^*mut/+*^ LP cells are associated with suppression of the NF-kB/SASP response.

**Fig. 4.**
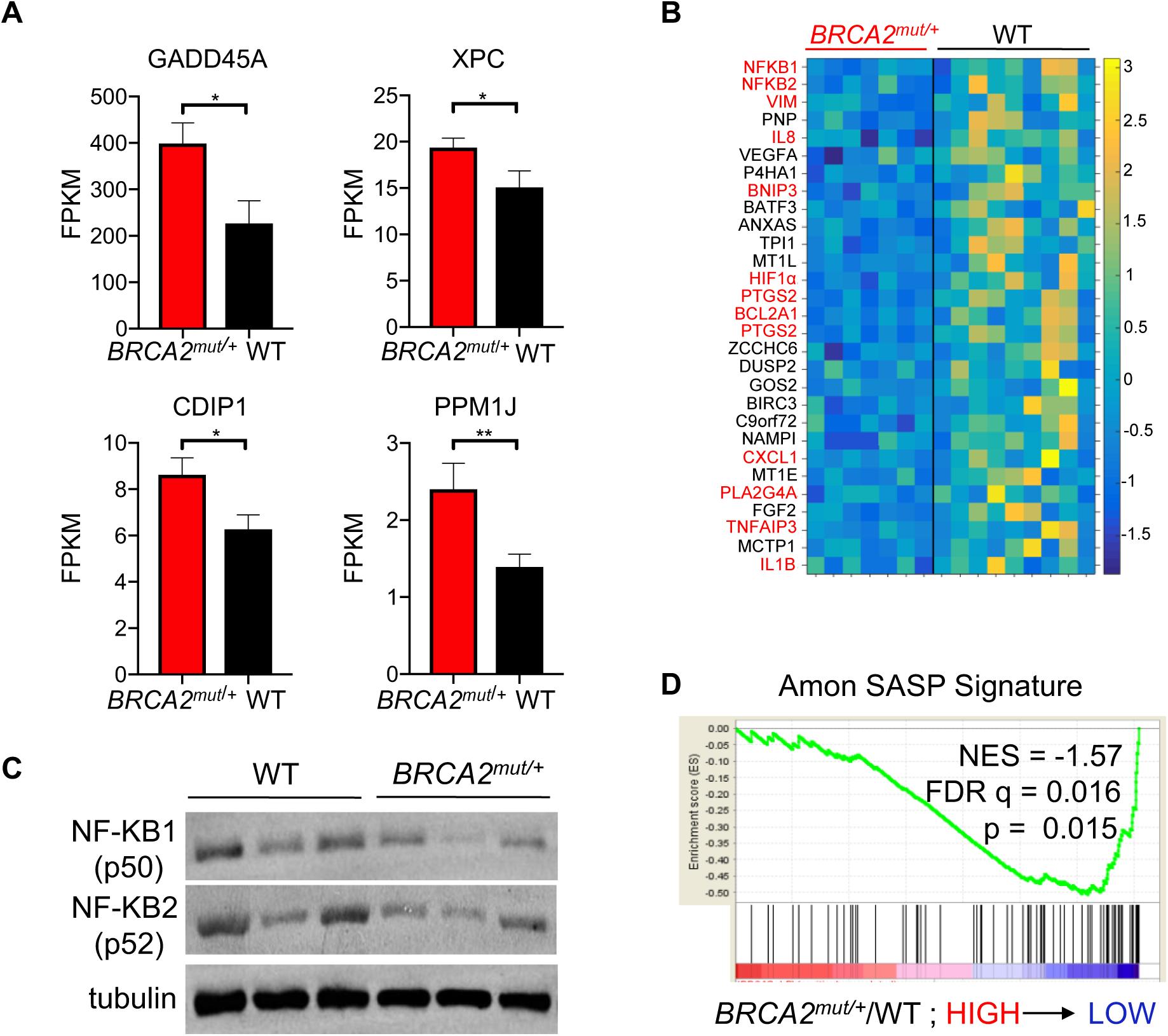
*BRCA2*^*mut/+*^ LP cells display increased TP53 activity and suppressed NF-kB/SASP pathway expression. **(A)** Bar charts show the mean expression levels of canonical TP53 target genes in freshly-sorted *BRCA2* carrier LP cells (n = 7 patients) compared to controls (WT, n = 9 patients), assessed by RNA-seq. Error bars denote standard error of the mean. p values by Mann-Whitney test. * p < 0.05, ** p < 0.01. **(B)** Heatmap depicts down-regulation of NF-kB/SASP pathway genes in *BRCA2* carrier LP cells compared to controls (WT), assessed by RNA-seq as in (A). Columns correspond to individual patients. Direct NF-kB target genes are highlighted in red. **(C)** Western blot analysis shows that NFKB1 (p50) and NFKB2 (p52) subunits are expressed at lower levels in *BRCA2*^*mut/+*^breast tissues compared to control (WT) tissues (n =3 patients per genotype). β-tubulin serve as loading control. **(D)** Negative enrichment of a SASP signature in GSEA analysis of RNA-seq data from freshly-sorted LP cells of *BRCA2* carriers (n = 7 patients) and controls (WT, n = 9 patients). NES, normalized enrichment score; FDR, false discovery rate.

### Age-associated deregulation of breast epithelial cell proportions in *BRCA2* carriers suggests expansion of a damaged LP cell population over time

Deregulated DNA damage and senescence/SASP responses in *BRCA2* LP cells might be expected to alter the proportion of these cells over time (*36*). We thus sought to address whether there were differences in the proportions of progenitor or other epithelial sub-populations in *BRCA2*^*mut/+*^ tissues compared to controls. We collected a larger cohort of tissues from *BRCA2* carriers (N=26) and controls (N=28), then performed flow cytometry analysis on these specimens and plotted the proportions of each epithelial sub-population as a function of age for each cohort (Fig. 5A). In non-carrier controls no significant age-associated changes in the prevalence of these sub-populations were noted. In contrast, *BRCA2* carriers showed an age-associated expansion in the proportion of LP cells and a decline in the basal cell fraction (Fig. 5B and fig. S5, A and B). These differences were not accounted for by demographic factors such as parity or menopausal status, as such factors were not associated with significant differences in epithelial cell proportions (fig. S5, C and D). Thus, DNA-damage and suppression of a senescence-associated program in *BRCA2*^*mut/+*^ LP cells is accompanied by an age-associated expansion of this progenitor cell compartment (*36*).

**Fig. 5.**
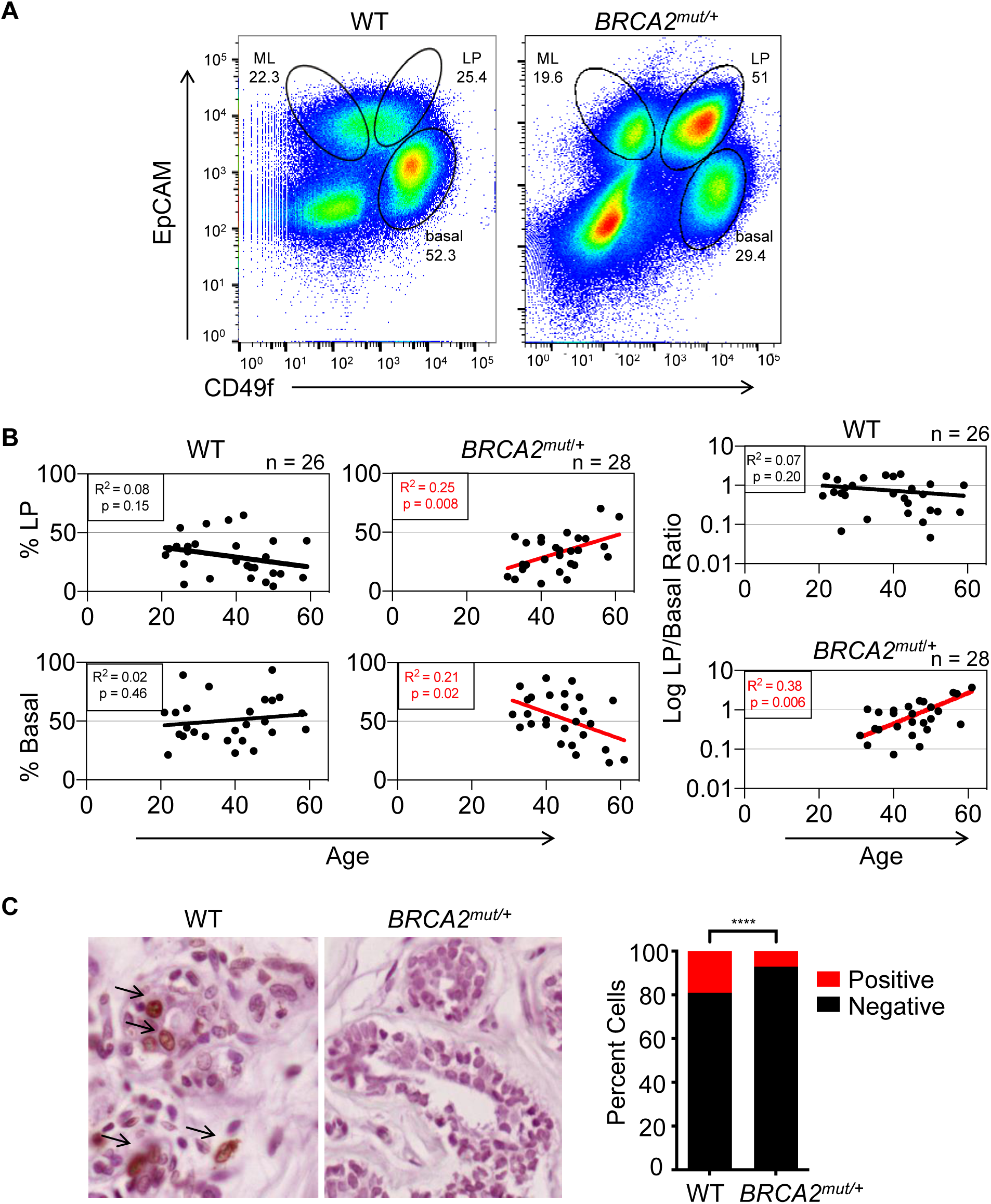
Non-cancerous breast tissues of *BRCA2* mutation carriers demonstrate age-associated deregulation of epithelial cell proportions compared to controls. **(A)** Representative flow cytometry analysis showing distinct epithelial subpopulations (basal, LP, mature luminal (ML)) isolated from breast tissues of control (WT) and *BRCA2* mutation carriers following sorting via CD49f and EpCAM staining. Numbers indicate percentages of each epithelial cell subpopulation. **(B)** Linear regression analysis of LP and basal cell proportions by age for controls (WT) (n = 26 patients) and *BRCA2* carriers (n = 28 patients). The LP/basal ratio by patient provides additional validation as it accounts for technical factors that may have subtle effects on absolute cell numbers. **(C)** TUNEL staining of representative control (WT) and *BRCA2* carrier tissues. Summary data obtained by counting four fields for 5 patients per genotype. ****p <0.0001 by Fisher’s exact test.

Finally, we hypothesized that altered epithelial cell proportions in *BRCA2* carrier tissues may be associated with differences in cell proliferation and/or survival in vivo. The relationship between NF-kB/SASP program, proliferation and susceptibility to apoptosis is a cell-type and context-dependent one (*36*). We did not observe strong differences in proliferation assessed by Ki67 staining between these *BRCA2* carrier breast tissues and controls (not shown), prompting us to ask whether differences in cell survival might contribute to the age-associated expansion of the LP population in this context. We therefore carried out TUNEL staining, an established marker of apoptosis, in *BRCA2* carrier tissues and controls. The proportion of TUNEL-positive cells is well-documented in normal human breast epithelial tissues, and we observed a similar prevalence of these cells in the control tissues we tested (Fig. 5C) (*37*). In contrast, however, *BRCA2* mutation carrier tissues consistently showed a paucity of TUNEL-positive luminal epithelial cells across all patients tested, in keeping with established links between checkpoint and NF-kB suppression and a defective apoptotic response (Fig. 5C) (*38, 39*). Collectively, our findings suggest that *BRCA2* haploinsufficiency is associated with an age-associated accumulation of DNA-damaged luminal epithelial progenitor cells bearing altered checkpoint and survival responses (fig. S6).

## Discussion

This study advances our understanding of early changes in *BRCA2*^*mut/+*^ breast tissues, defining unanticipated phenotypes in this setting with implications for both cancer risk assessment and prevention. The majority of the tissues we studied were deemed to be histologically normal by highly experienced breast pathologists, suggesting that the alterations we report precede clinically-defined cellular abnormalities (tables S1, S2). We present evidence that a failed replication stress response and DNA damage in *BRCA2*^*mut/+*^ tissues results from haploinsufficiency for *BRCA2* rather than homozygous loss of function. While the presence of haploinsufficiency for either *BRCA1* or *BRCA2* in vivo has been controversial, our findings are in accord with data suggesting that LOH for the wild-type *BRCA2* is not universal in *BRCA2*-associated cancers (*40*). Our observations are also in keeping with a recent report that the BRCA2 protein is selectively susceptible to degradation by environmental aldehydes (*41*), an effect which could contribute to a haploinsufficient phenotype in cells with only one functional *BRCA2* allele. Nonetheless, our study of early pathogenesis does not prove that haploinsufficiency is sufficient for progression to invasive malignancy, but rather we define a haploinsufficient phenotype for *BRCA2* that is a potential initiating event for these cancers.

A prominent feature of the phenotype we have uncovered is frequent sub-chromosomal aneuploidy, most prevalent within the LP cell population. LP cells are a potential target cell for *BRCA2*-associated carcinogenesis in the breast, and indeed we observe instances of apparently clonally related genomic alterations among these cells. Such alterations could conceivably represent the earliest somatic genetic abnormalities that underlie these malignancies. Notably, all the CNVs we identified were sub-chromosomal and therefore are to be distinguished from whole-chromosome gains and losses that are typically later events and are associated with TP53 inactivation (*42*). Although these early genomic changes are likely to include many passenger events, they nevertheless may provide a quantifiable hallmark of the pre-neoplastic *BRCA2* carrier state. Tracking the prevalence of DNA-damaged cells in the clinical setting could possibly improve risk prediction for such women, who are faced with the difficult choice of whether to undergo mastectomy long before cancer develops. Finally, the *BRCA2* haploinsufficient phenotype we report may portend particular vulnerabilities of certain *BRCA2*^*mut/+*^ cancer precursor cells. Accordingly, this work provides a foundation for future studies seeking to identify novel pharmacologic approaches to cancer prevention in this setting.

## Materials and Methods

### Human breast tissues

Fresh human breast tissues were obtained from Massachusetts General Hospital with approval by the local Institution Review Board and signed informed patient consent (Protocols 93-085 and 2008-P-1789). Samples were either normal breast tissues from reduction mammoplasties (confirmed by pathology) or non-cancerous breast tissues from prophylactic mastectomies of known BRCA1 or BRCA2 mutation carriers. All BRCA1/2 carrier status was determined through clinical germline genetic testing performed by commercial providers prior to tissue collection.

### Mammary cell preparations

Tissue samples were minced and digested with collagenase/hyaluronidase (Stemcell technologies) in complete Epicult-B Medium supplemented with 0.48 µg/mL hydrocortisone (Stemcell technologies) overnight at 37 °C. The resulting suspensions were either cryopreserved or further sequentially digested with 0.25% trypsin, 5 mg/mL dispase and 1 mg/mL DNase I. Single cell suspensions were collected by filtration through a 40 µm cell strainer.

### Cell staining and sorting

Cells were blocked with rat immunoglobulin (Jackson Immunolabs) and antibody to Fc receptor binding inhibitor (eBioscience) before incubation with the following primary antibodies: PE-conjugated anti-human CD31 (BD Pharmingen), PE-conjugated anti-human CD45 (BD Pharmingen), PE-conjugated anti-human CD235a (BD Pharmingen), BV650-conjugated anti-human EPCAM CD326 (Biolegend), biotin-conjugated anti-human ITGA6 (eBioscience). Where required, cells were incubated with APC-Cy7-conjugated streptavidin (BD Pharmingen). Cells were either stained with DAPI for viability or fixed with 1% PFA and stained with zombie aqua fixable viability kit (Biolegend). Viable cells were sorted on a FACSAria flow cytometer (Becton Dickinson). Data was analyzed using FlowJo software (Tree Star).

### Single-Cell PCR for allele-specific LOH analysis

Microaspirated single cells were transferred into PCR tubes containing lysis buffer (water + 400 ng/ul Proteinase K + 17 µM SDS) and DNA was amplified by nested PCR using primers flanking BRCA2 mutations. Sanger sequencing was performed by the CCIB DNA Core Facility at Massachusetts General Hospital.

Primer sequences for patient 128 (Val3079PhefsX4) are as in the following:

1^st^ PCR-Forward: TGGCGTCCATCATCAGATTT

1^st^ PCR-Reverse: TCAGAGGTTCAAAGAGGCTTAC

2^nd^ PCR-Forward: CAGATTTACCAGCCACGGGA

2^nd^ PCR-Reverse: GCCAACTGGTAGCTCCAACTAA

Primer sequences for patient 140 (6027del4) are as in the following:

1^st^ PCR-Forward: GGGCCACCTGCATTTAGGAT

1^st^ PCR-Reverse: TGAGCTGGTCTGAATGTTCGT

2^nd^ PCR-Forward: GCAGGTTGTTACGAGGCATT

2^nd^ PCR-Reverse: CCTGGACAGATTTTCCACTTGC

### Comet Assays

Single cell suspensions from patient samples were plated in ultralow-adherence plates in DMEM/F12 medium containing 5 µg/ml insulin, 10 ng/mL EGF, 5 ng/ml bFGF, 4 µg/ml heparin, 500 ng/ml hydrocortisone, B27, Glutamax and penicillin-streptomycin. Cells were either treated with hydroxyurea (10 mM, Sigma) for 4 h or left untreated, washed with PBS and alkaline comet assays were performed using Trevigen Comet Assay kit, according to the manufacturer’s instructions. Olive tail movement was quantified with ImageJ, 50 individual cells were quantified per condition.

### Immunostaining

Immunofluorescence for paraffin sections and TUNEL staining were performed by Dana-Farber/Harvard Cancer Center Specialized Histopathology Core. For Immunofluorescence in cells, fixation was performed with methanol for 10 min followed by permeabilization in 0.1% TritonX100 for 2 min. Blocking was performed with 10% horse serum for 30 min and cells were further incubated with primary p-CHK1 antibody (Novus Biologicals) for 2 hr, washed with wash buffer (PBS +10% horse serum + 0.1% TritonX100), incubated with appropriate secondary antibody for 1hr and stained with DAPI. All immunofluorescence images were captured by a confocal microscope (Leica TCS SP8) and were analyzed by ImageJ.

### Western Blotting

Snap-frozen tissues were homogenized using Precellys 24 homogenizer (Bertin Technologies). For total protein extraction, cells were lysed in RIPA buffer (10 mM Tris-HCl pH 7.5, 150 mM NaCl, 1 mM EDTA, 1% sodium deoxycholate, 0.1% (w/v) SDS, 1% (v/v) NP40, proteinase inhibitor cocktail, phosphatase inhibitor cocktail) for 30 min on ice. Western blotting was performed using NFKB p50 (Santa Cruz Biotechnologies) and NFKB p52 (Millipore) antibodies by standard protocol.

### Single-Cell Copy Number Analysis

Fresh tissues were dissociated as described above and single cells were isolated by microaspiration. Genomic DNA was amplified and sequenced as described in Knouse et al. 2014. Fastqs were aligned using bwa-mem, with resulting bams sorted and duplicates marked using Picard. Coverage was then computed over 500kb bins across the entire genome. The count for each bin was then divided by the sum across all bins for the relative sample (to correct for library size), and then by the median for that genomic bin across all samples from the same batch. The coverage profiles were then transformed into .wig files and fed into the R package HMMCopy for segmentation and CNV-calling. HMMCopy was run with e = 0.9999999 and nu = 5, with all other parameters set to default. A noise statistic termed “VS” was computed in the same manner as Knouse et al. 2014, with cells with values greater than or equal to 0.5 being excluded from the analysis. CNVs that mapped to the Y chromosome, were less than 10Mb in size, or had an absolute log2 ratio less than 0.4 were excluded from the analysis.

### RNA-Seq Analysis

Total RNA from sorted cell populations was extracted using RNeasy FFPE kit, according to manufacturer’s instructions. Libraries for ribosomally reduced RNA was prepared by Harvard Biopolymers Facility using directional RNA-seq Wafergen protocol. Libraries were sequenced on Illumina HiSeq 2000 at Next Generation Sequencing Core at Massachusetts General Hospital. TPM values were computed using Salmon and batch-corrected using ComBat. The two samples with the lowest total counts were excluded from the analysis. GSEA was run on the ComBat-corrected TPM values using phenotype permutation and default parameters. The heatmaps in figures 4B and 4D were made using the ComBat-corrected TPM values, subset to the comparison of interest, and transformed into z-scores by gene.

### Other Statistical Methods

p Values were determined using the Student’s unpaired t test unless indicated otherwise.

## Supplementary Materials

### Materials and methods

Fig. S1. Analysis of single-cell WGS data from normal human skin and brain cells.

Fig. S2. Identification and characterization of CNVs in freshly collected *BRCA2*^*mut/+*^ breast epithelial cells.

Fig. S3. Characterization of replication stress response deficiency and haploinsufficiency in

*BRCA2*^*mut/+*^ breast epithelial cells.

Fig. S4. Suppression of NF-kB/SASP response associated with loss of *BRCA2*.

Fig. S5. Proportions of mammary epithelial cell subsets in *BRCA2* carrier and control tissues.

Fig. S6. Summary of findings reflecting haploinsufficiency in primary *BRCA2*^*mut/+*^ breast epithelial cells.

Table S1. Characteristics of patients undergoing whole-genome sequencing of breast tissues.

Table S2. Characteristics of patients undergoing RNA-sequencing of breast tissues.

## Acknowledgements

We thank the HSCI-CRM Flow Cytometry Core Facility for assistance with cell sorting and MGH DF/HCC Specialized Histopathology Service Core for immunostaining experiments. We thank Biopolymers Facility at Harvard Medical School for library processing of RNA samples and MGH Next Generation Sequencing Core for performing RNA-sequencing. We are grateful to MIT BioMicro Center for performing genome sequencing reactions.

## Funding

This work was supported by DOD/CDMRP grant BC140903 (L.W.E.), by the Tracey Davis Breast Cancer Research Fund (L.W.E.), by the Weissman Family MGH Research Scholar grant (L.W.E.), by the Susan G. Komen Foundation grant PDF16380794 (M.K.), by a Terri Brodeur Breast Cancer Foundation grant (M.K.), and by the Howard Hughes Medical Institute (A.A.).

## Author Contributions

M. K., A.A. and L.W.E. conceived and designed the study. M.C.S. contributed patient samples. M.K., R.E.S. and S.V.S. designed and performed experiments and interpreted data. H.R. and R.M. and V.V. performed experiments. M.K., E.Z., A.D., M.Y. performed data analysis. M.K, A.L., K.R., S.R. and M.L. performed bioinformatic analysis and interpreted data. L.W.E., A.A. and M.L. conceived experiments, interpreted data and provided funding. L.W.E. and M.K. wrote the manuscript. All authors approved the final submitted manuscript.

The authors declare no competing financial interests.

## References

1. H. Davies et al., HRDetect is a predictor of BRCA1 and BRCA2 deficiency based on mutational signatures. Nat Med 23, 517–525 (2017).

2. F. J. Couch, K. L. Nathanson, K. Offit, Two decades after BRCA: setting paradigms in personalized cancer care and prevention. Science 343, 1466–1470 (2014).

3. T. Ludwig, D. L. Chapman, V. E. Papaioannou, A. Efstratiadis, Targeted mutations of breast cancer susceptibility gene homologs in mice: lethal phenotypes of Brca1, Brca2, Brca1/Brca2, Brca1/p53, and Brca2/p53 nullizygous embryos. Genes Dev 11, 1226–1241 (1997).

4. S. K. Sharan et al., Embryonic lethality and radiation hypersensitivity mediated by Rad51 in mice lacking Brca2. Nature 386, 804–810 (1997).

5. A. Suzuki et al., Brca2 is required for embryonic cellular proliferation in the mouse. Genes Dev 11, 1242–1252 (1997).

6. A. Tutt et al., Cell cycle and genetic background dependence of the effect of loss of BRCA2 on ionizing radiation sensitivity. Oncogene 22, 2926–2931 (2003).

7. W. Feng, M. Jasin, BRCA2 suppresses replication stress-induced mitotic and G1 abnormalities through homologous recombination. Nat Commun 8, 525 (2017).

8. F. Connor et al., Tumorigenesis and a DNA repair defect in mice with a truncating Brca2 mutation. Nat Genet 17, 423–430 (1997).

9. L. C. Gowen, B. L. Johnson, A. M. Latour, K. K. Sulik, B. H. Koller, Brca1 deficiency results in early embryonic lethality characterized by neuroepithelial abnormalities. Nat Genet 12, 191–194 (1996).

10. R. Hakem et al., The tumor suppressor gene Brca1 is required for embryonic cellular proliferation in the mouse. Cell 85, 1009–1023 (1996).

11. Pathology of familial breast cancer: differences between breast cancers in carriers of BRCA1 or BRCA2 mutations and sporadic cases. Breast Cancer Linkage Consortium. Lancet 349, 1505–1510 (1997).

12. G. Molyneux et al., BRCA1 basal-like breast cancers originate from luminal epithelial progenitors and not from basal stem cells. Cell Stem Cell 7, 403–417 (2010).

13. E. Lim et al., Aberrant luminal progenitors as the candidate target population for basal tumor development in BRCA1 mutation carriers. Nat Med 15, 907–913 (2009).

14. J. E. Visvader, J. Stingl, Mammary stem cells and the differentiation hierarchy: current status and perspectives. Genes Dev 28, 1143–1158 (2014).

15. K. A. Knouse, J. Wu, A. Amon, Assessment of megabase-scale somatic copy number variation using single-cell sequencing. Genome Res 26, 376–384 (2016).

16. K. A. Knouse, J. Wu, C. A. Whittaker, A. Amon, Single cell sequencing reveals low levels of aneuploidy across mammalian tissues. Proc Natl Acad Sci U S A 111, 13409–13414 (2014).

17. N. Cancer Genome Atlas, Comprehensive molecular portraits of human breast tumours. Nature 490, 61–70 (2012).

18. S. Nik-Zainal et al., Landscape of somatic mutations in 560 breast cancer whole-genome sequences. Nature 534, 47–54 (2016).

19. X. Song et al., alpha-MSH activates immediate defense responses to UV-induced oxidative stress in human melanocytes. Pigment Cell Melanoma Res 22, 809–818 (2009).

20. E. Nolan et al., RANK ligand as a potential target for breast cancer prevention in BRCA1-mutation carriers. Nat Med 22, 933–939 (2016).

21. W. Feng, M. Jasin, Homologous Recombination and Replication Fork Protection: BRCA2 and More! Cold Spring Harb Symp Quant Biol 82, 329–338 (2017).

22. N. Walworth, S. Davey, D. Beach, Fission yeast chk1 protein kinase links the rad checkpoint pathway to cdc2. Nature 363, 368–371 (1993).

23. J. Bartek, J. Lukas, Chk1 and Chk2 kinases in checkpoint control and cancer. Cancer Cell 3, 421–429 (2003).

24. D. J. McGrail et al., Defective Replication Stress Response Is Inherently Linked to the Cancer Stem Cell Phenotype. Cell Rep 23, 2095–2106 (2018).

25. T. Mai et al., Estrogen receptors bind to and activate the HOXC4/HoxC4 promoter to potentiate HoxC4-mediated activation-induced cytosine deaminase induction, immunoglobulin class switch DNA recombination, and somatic hypermutation. J Biol Chem 285, 37797–37810 (2010).

26. N. M. Robert, J. J. Tremblay, R. S. Viger, Friend of GATA (FOG)-1 and FOG-2 differentially repress the GATA-dependent activity of multiple gonadal promoters. Endocrinology 143, 3963–3973 (2002).

27. S. A. Ghandhi, B. Yaghoubian, S. A. Amundson, Global gene expression analyses of bystander and alpha particle irradiated normal human lung fibroblasts: synchronous and differential responses. BMC Med Genomics 1, 63 (2008).

28. J. Jonkers et al., Synergistic tumor suppressor activity of BRCA2 and p53 in a conditional mouse model for breast cancer. Nat Genet 29, 418–425 (2001).

29. J. Smith, L. M. Tho, N. Xu, D. A. Gillespie, The ATM-Chk2 and ATR-Chk1 pathways in DNA damage signaling and cancer. Adv Cancer Res 108, 73–112 (2010).

30. M. T. M. van Jaarsveld, D. Deng, E. A. C. Wiemer, Z. Zi, Tissue-Specific Chk1 Activation Determines Apoptosis by Regulating the Balance of p53 and p21. iScience 12, 27–40 (2019).

31. S. S. McDade et al., Genome-wide characterization reveals complex interplay between TP53 and TP63 in response to genotoxic stress. Nucleic Acids Res 42, 6270–6285 (2014).

32. S. T. Younger, D. Kenzelmann-Broz, H. Jung, L. D. Attardi, J. L. Rinn, Integrative genomic analysis reveals widespread enhancer regulation by p53 in response to DNA damage. Nucleic Acids Res 43, 4447–4462 (2015).

33. J. P. Coppe et al., Senescence-associated secretory phenotypes reveal cell-nonautonomous functions of oncogenic RAS and the p53 tumor suppressor. PLoS Biol 6, 2853–2868 (2008).

34. C. D. Wiley et al., Small-molecule MDM2 antagonists attenuate the senescence-associated secretory phenotype. Sci Rep 8, 2410 (2018).

35. C. J. Sieben, I. Sturmlechner, B. van de Sluis, J. M. van Deursen, Two-Step Senescence-Focused Cancer Therapies. Trends Cell Biol 28, 723–737 (2018).

36. B. G. Childs, D. J. Baker, J. L. Kirkland, J. Campisi, J. M. van Deursen, Senescence and apoptosis: dueling or complementary cell fates? EMBO Rep 15, 1139–1153 (2014).

37. F. Feuerhake, W. Sigg, E. A. Hofter, P. Unterberger, U. Welsch, Cell proliferation, apoptosis, and expression of Bcl-2 and Bax in non-lactating human breast epithelium in relation to the menstrual cycle and reproductive history. Breast Cancer Res Treat 77, 37–48 (2003).

38. M. B. Kastan, J. Bartek, Cell-cycle checkpoints and cancer. Nature 432, 316–323 (2004).

39. S. J. Veuger, B. W. Durkacz, Persistence of unrepaired DNA double strand breaks caused by inhibition of ATM does not lead to radio-sensitisation in the absence of NF-kappaB activation. DNA Repair (Amst) 10, 235–244 (2011).

40. K. N. Maxwell et al., BRCA locus-specific loss of heterozygosity in germline BRCA1 and BRCA2 carriers. Nat Commun 8, 319 (2017).

41. S. L. W. Tan et al., A Class of Environmental and Endogenous Toxins Induces BRCA2 Haploinsufficiency and Genome Instability. Cell 169, 1105–1118 e1115 (2017).

42. L. Sansregret, C. Swanton, The Role of Aneuploidy in Cancer Evolution. Cold Spring Harb Perspect Med 7, (2017).

43. M. R. Ramsey et al., FGFR2 signaling underlies p63 oncogenic function in squamous cell carcinoma. J Clin Invest 123, 3525–3538 (2013).

