## supplementary material for "Aneuploidy and deregulated DNA damage response define haploinsufficiency in breast tissues of *BRCA2* mutation carriers"

Mihriban Karaayvaz^1^, Rebecca E Silberman^2#^, Adam Langenbucher^1#^, Srinivas Vinod Saladi^1,3#^, Kenneth N Ross^1,4^, Elena Zarcaro^1^, Andrea Desmond^1^, Murat Yildirim^5^, Varunika Vivekanandan^1^, Hiranmayi Ravichandran^6^, Ravindra Mylavagnanam^6^, Michelle C Specht^7^, Sridhar Ramaswamy^1,4^, Michael Lawrence^1,4^, Angelika Amon^2^, Leif W Ellisen^1*^ 

^*^Correspondence:	Leif W. Ellisen, M.D., Ph.D.

MGH Cancer Center

CPZN-4204, 185 Cambridge Street

Boston, MA 02114

**Includes:**

- **Materials and methods**
- **Fig. S1. Analysis of single-cell WGS data from normal human skin and brain cells.**
- **Fig. S2. Identification and characterization of CNVs in freshly collected *BRCA2^mut/+^* breast epithelial cells.**
- **Fig. S3. Characterization of replication stress response deficiency and haploinsufficiency in *BRCA2^mut/+^* breast epithelial cells.**
- **Fig. S4. Suppression of NF-kB/SASP response associated with loss of *BRCA2*.**
- **Fig. S5. Proportions of mammary epithelial cell subsets in *BRCA2* carrier and control tissues.**
- **Fig. S6. Summary of findings reflecting haploinsufficiency in primary *BRCA2^mut/+^* breast epithelial cells.**
- **Table S1. Characteristics of patients undergoing whole-genome sequencing of breast tissues.**
- **Table S2. Characteristics of patients undergoing RNA-sequencing of breast tissues.**

**Materials and methods**

**Lentiviral BRCA2 knockdown and gene expression analysis**

BRCA2 knockdown in non-transformed MCF10A mammary epithelial cells was performed using lentiviral shRNA. The corresponding shRNA sequences for BRCA2 are GCAGCCATTAAATTGTCCATA and GCCTTGAATAATCACAGGCAA. Production of virus was performed as described (*43*). Following brief passage, total RNA was isolated by STAT60 (Tel-Test Inc, No. CS-111) following the manufacturer’s instructions. Gene expression analysis was performed by qRT-PCR (1 µg total RNA as template) using SuperScript IV (Invitrogen, No. 18091050) and iTaq Universal SYBR Green Supermix (Biorad, No.1725121) following the manufacturer’s instructions.

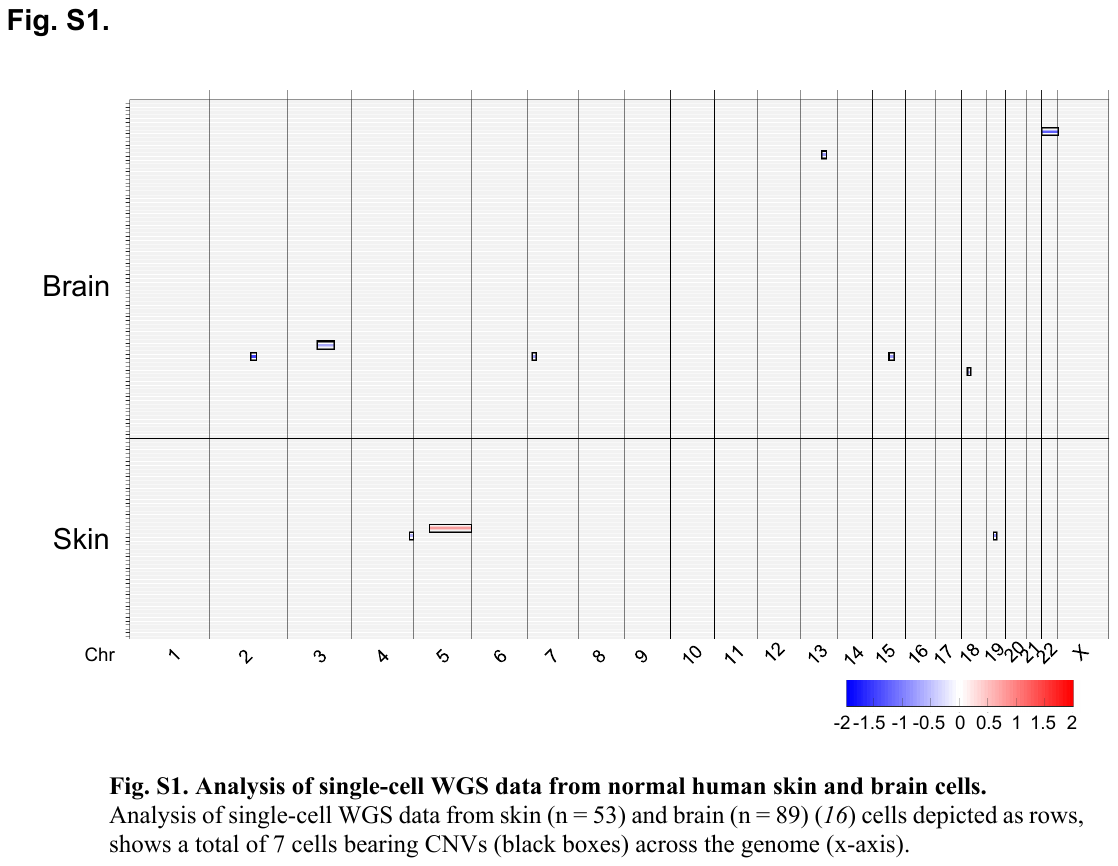

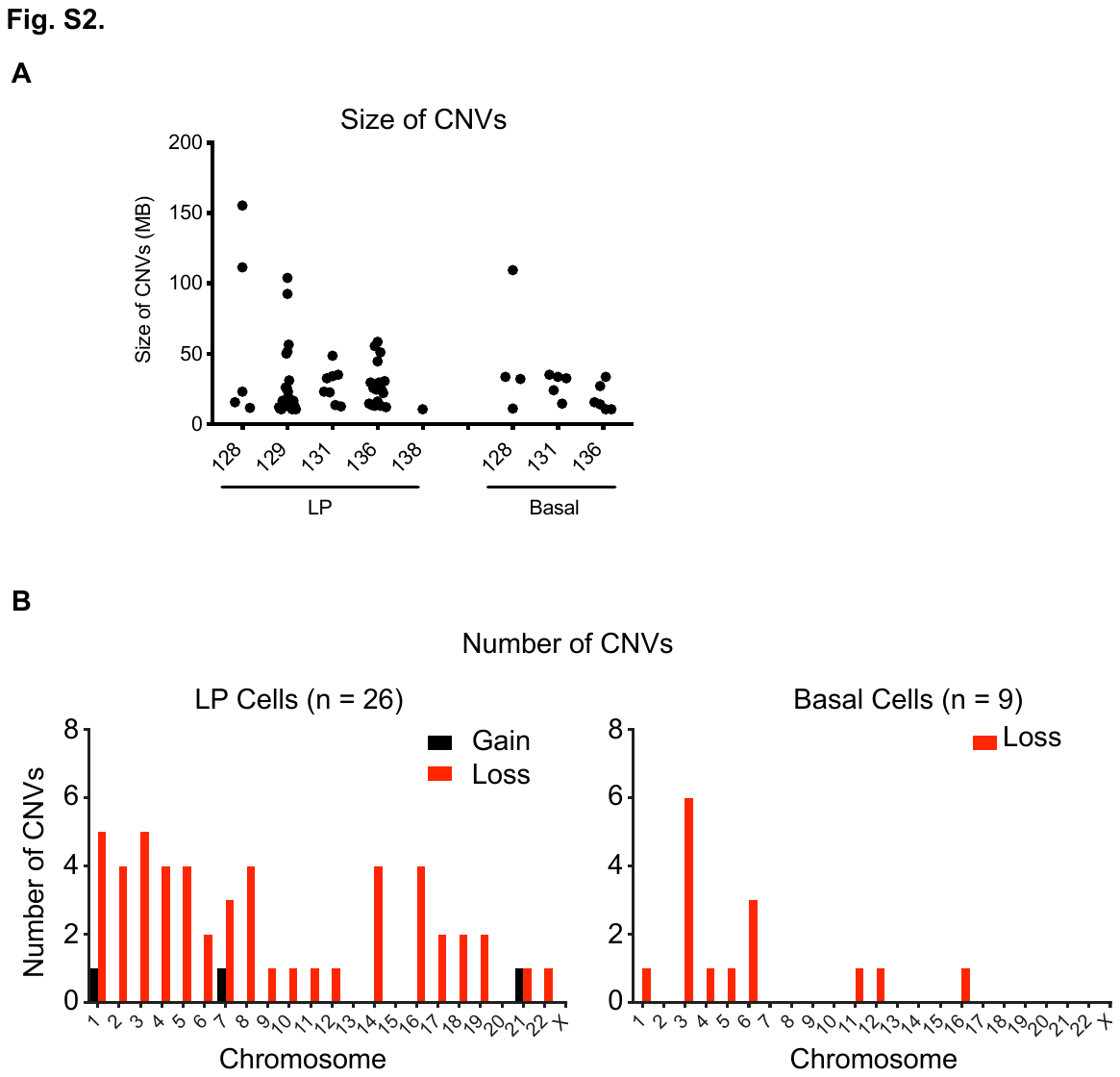

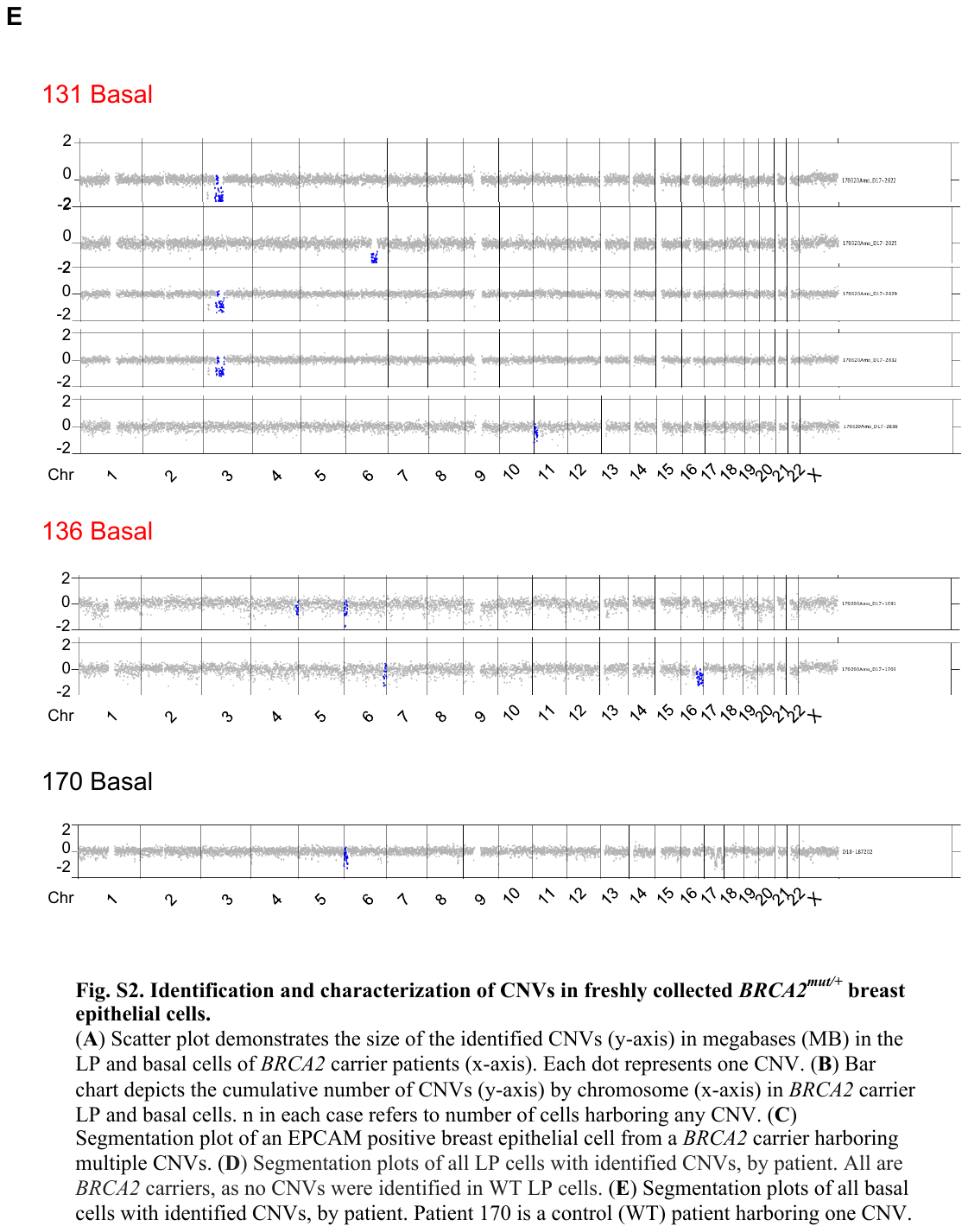

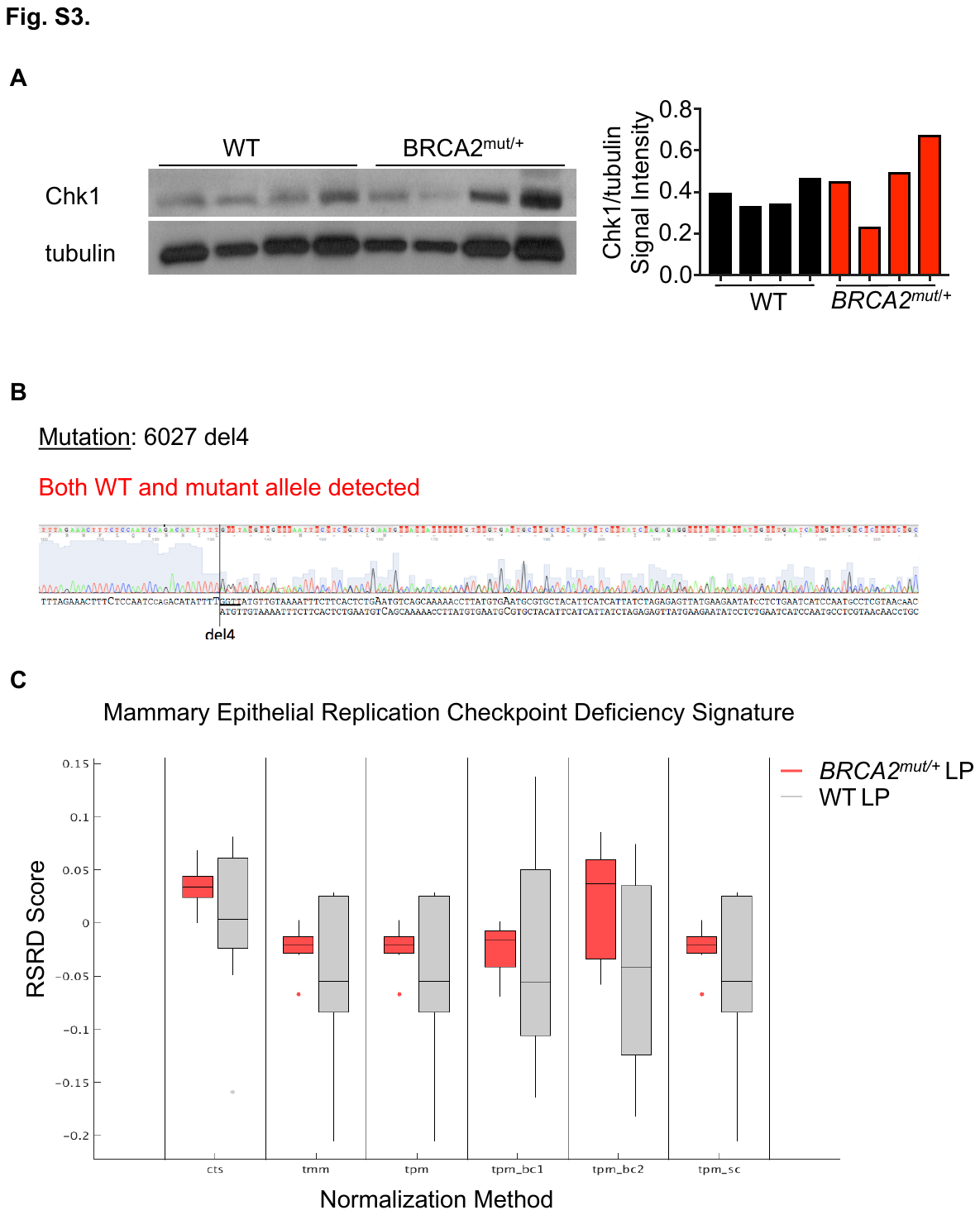

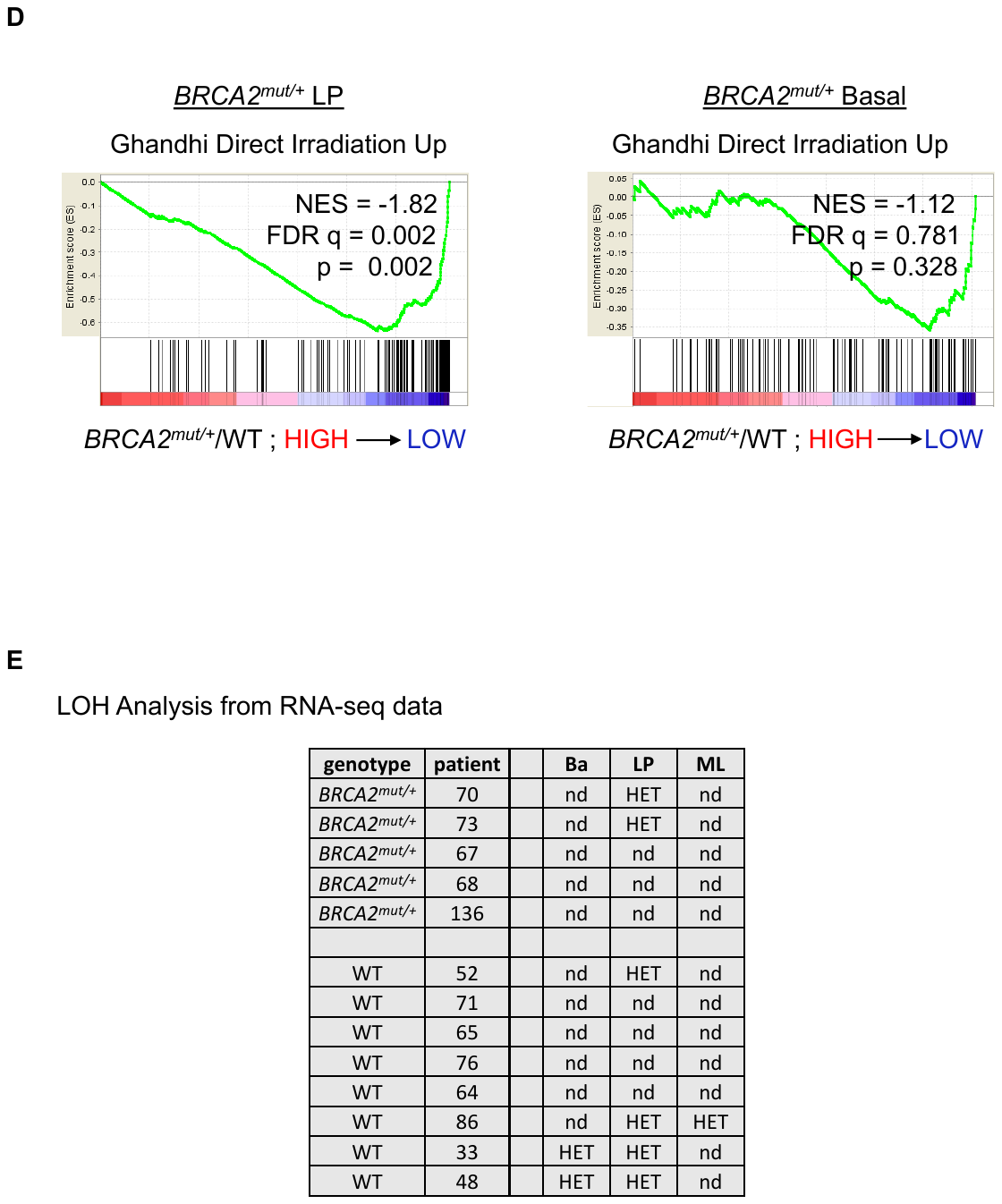

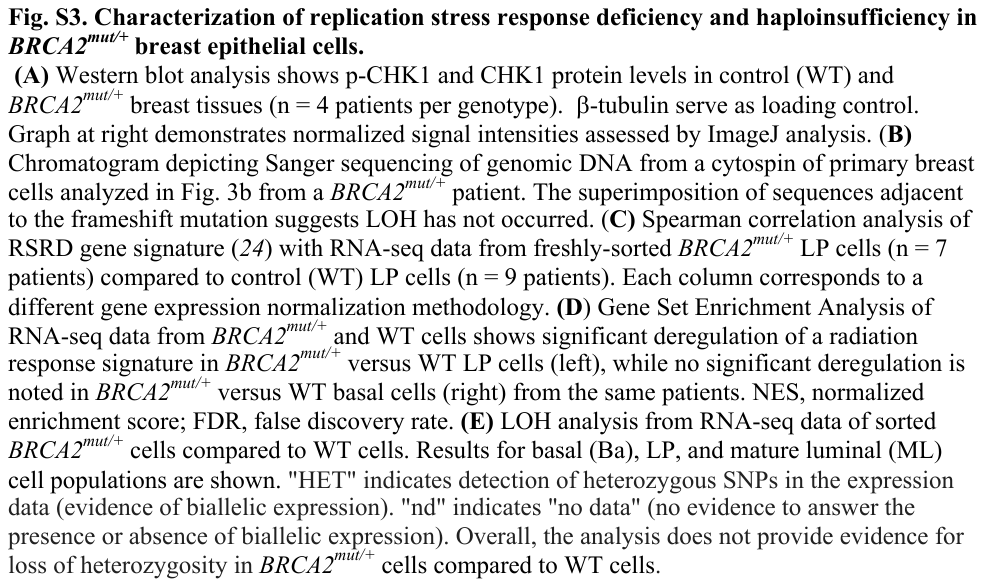

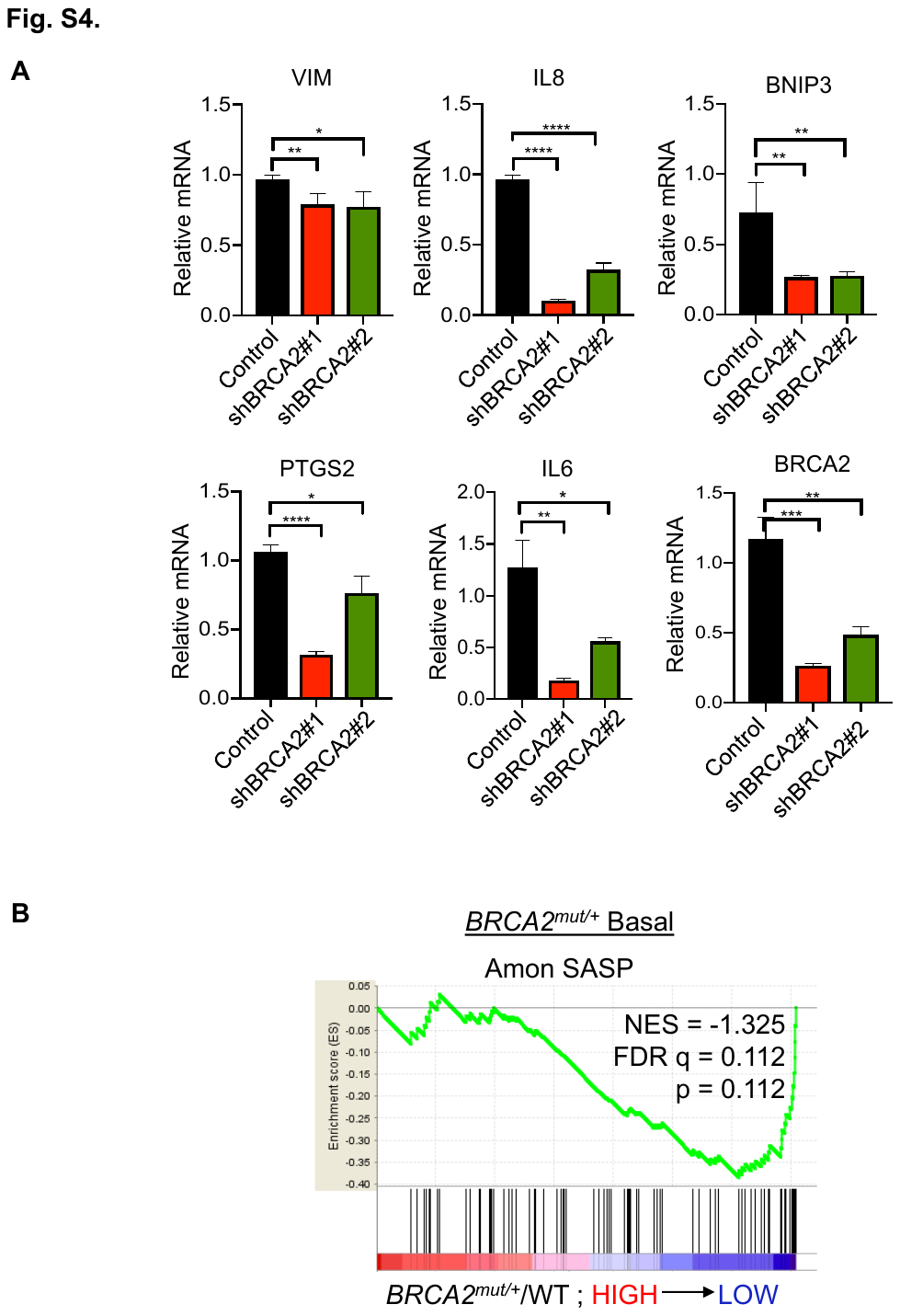

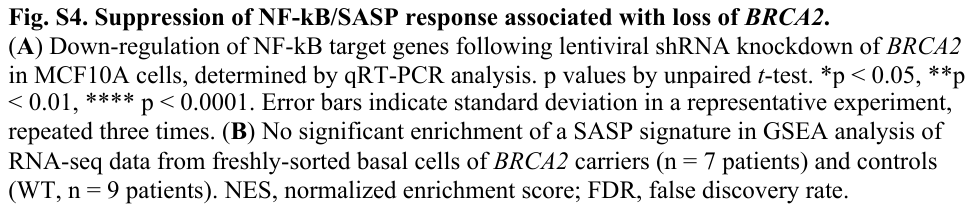

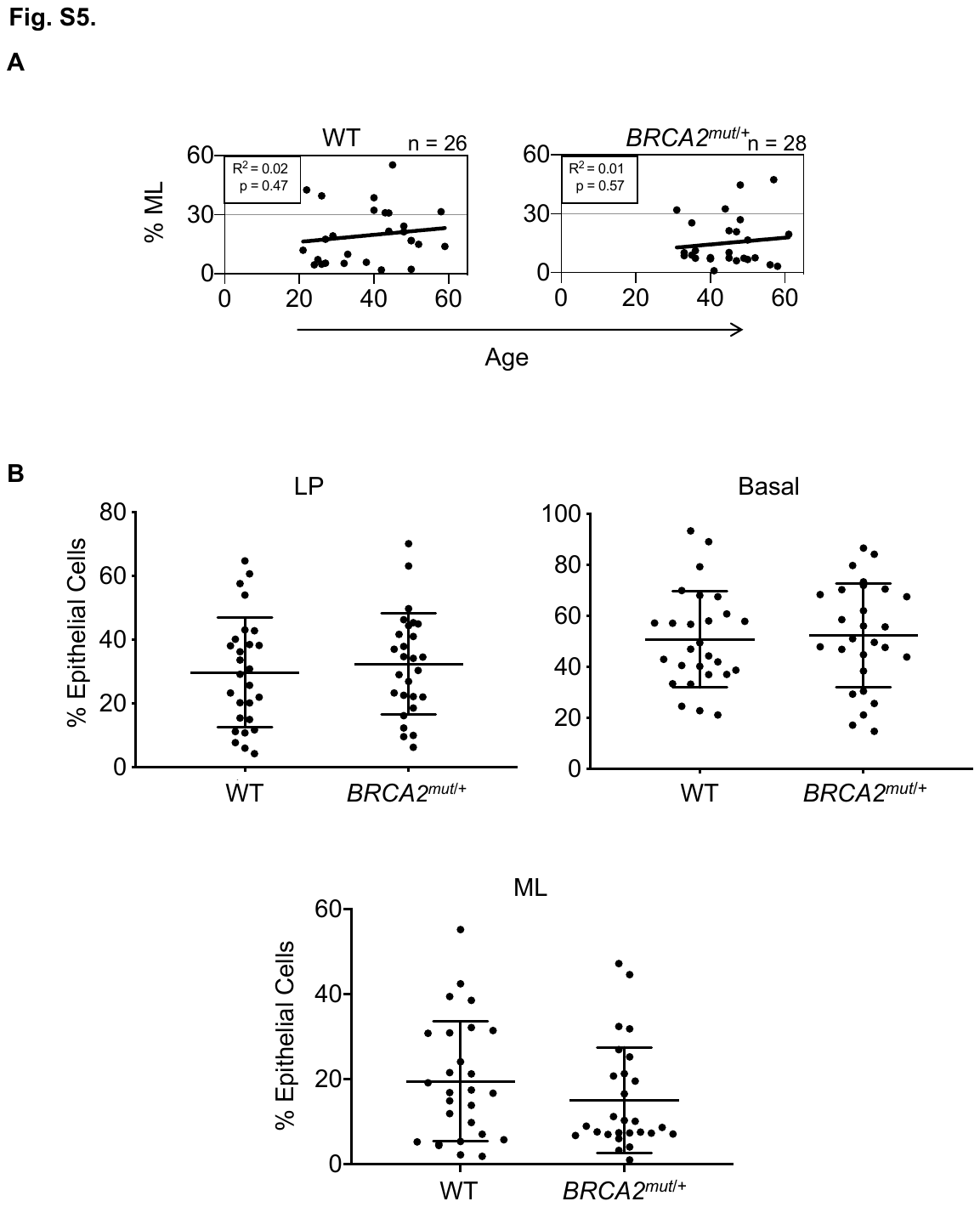

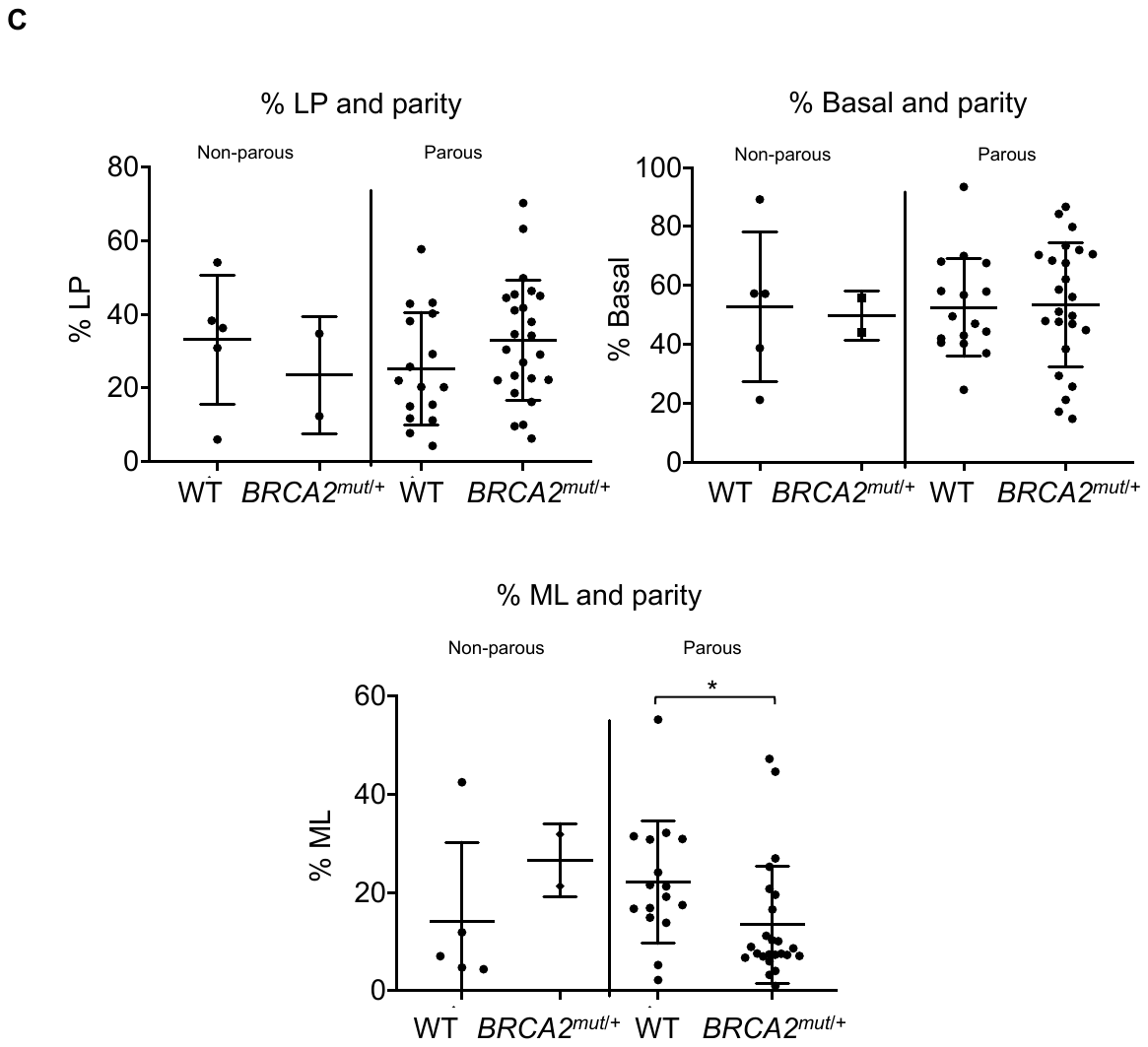

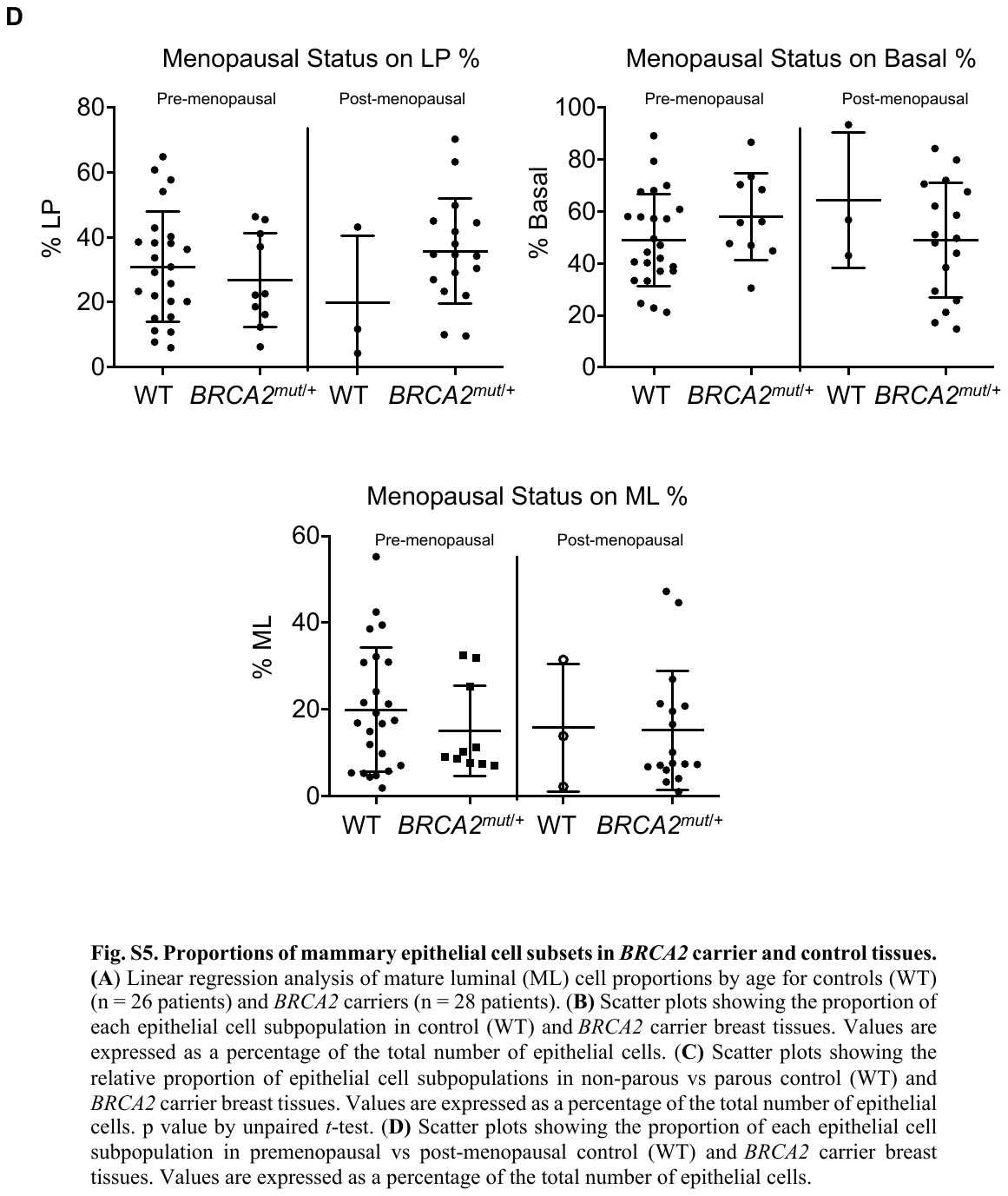

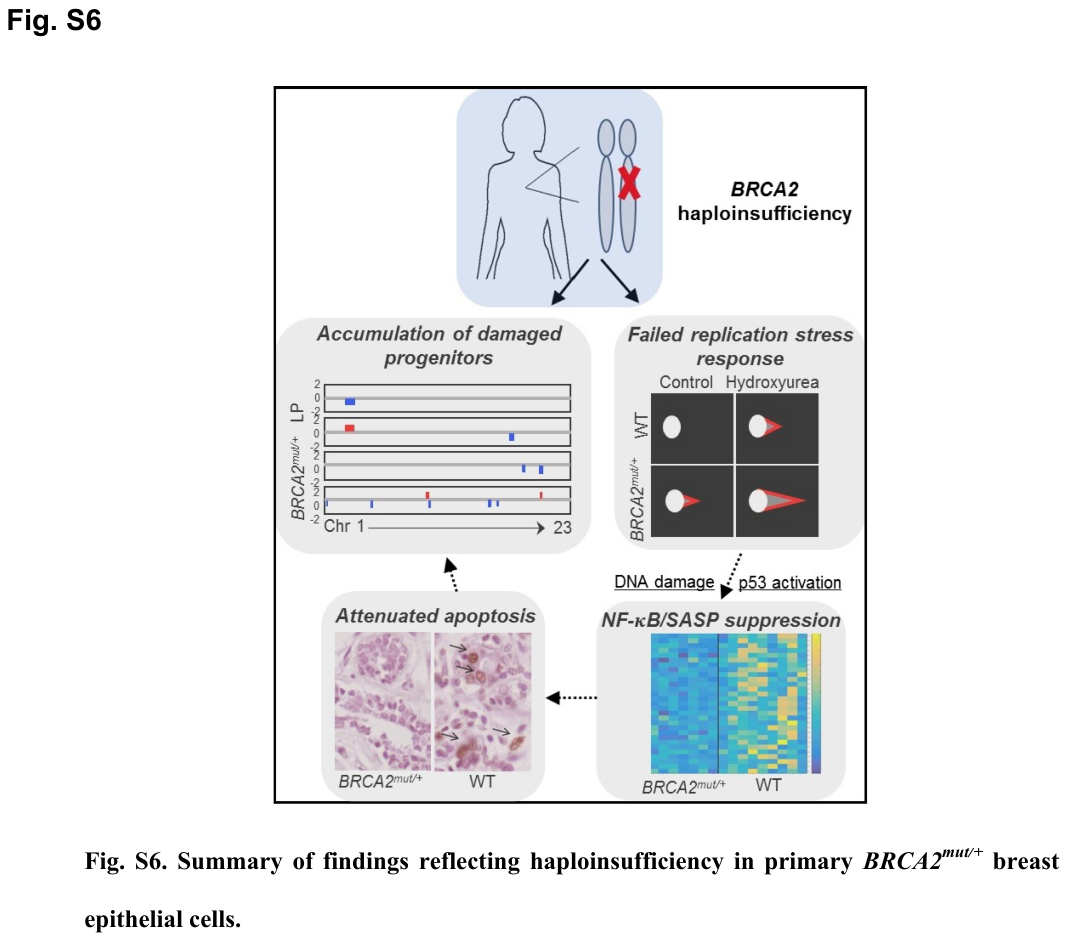

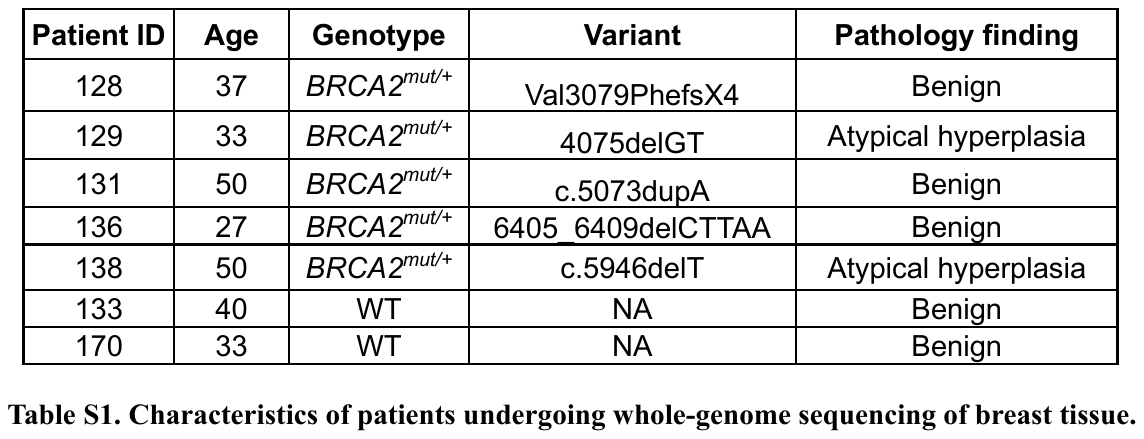
**Supplementary tables**

**Table S1.**

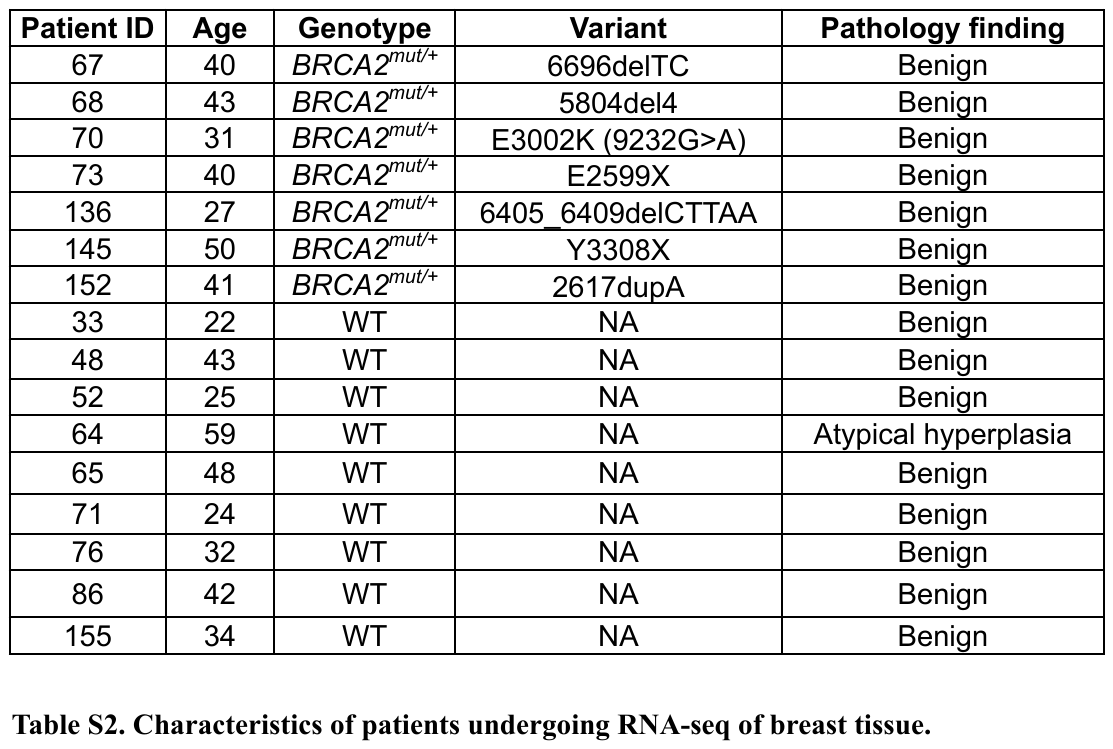
**Table S2.**
